# Neonatal Fc receptor is a functional receptor for human astrovirus

**DOI:** 10.1101/2022.11.13.516297

**Authors:** Kei Haga, Reiko Takai-Todaka, Akiko Kato, Akira Nakanishi, Kazuhiko Katayama

## Abstract

Human astrovirus (HAstV) is a global cause of gastroenteritis in infants, the elderly, and immunocompromised people. However, its infection mechanism is not fully understood, with its functional receptor not yet discovered. Here, we identify neonatal Fc receptor (FcRn) as a functional receptor for HAstV (mamastrovirus 1) using genome-wide CRISPR-Cas9 library screening in Caco2 cells. Deletion of *FCGRT* or *B2M*, which encode subunits of FcRn, rendered Caco2 cells and intestinal organoid cells unsusceptible to HAstV. We also show that human FcRn expression renders non-permissive MDCK cells susceptible and that FcRn directly binds HAstV spike protein. Thus, our findings provide insight into the entry mechanism of HAstV.

## Introduction

Astrovirus (family *Astroviridae*) is a nonenveloped virus with a positive single-stranded RNA genome that causes gastroenteritis in humans and animals worldwide. Astroviruses are classified by their capsid amino acid sequences into avastroviruses and mamastroviruses, which infect birds and mammals, respectively. Mamastroviruses comprise 19 groups and include the human-infecting mamastroviruses 1 (HAstV serotypes 1–8), 6 (MLB1 and 2), 8 (VA2, 4, and 5), and 9 (VA1 and 3) ^1^.

Classical-type human astroviruses (e.g., mamastrovirus 1), unlike novel-type astroviruses (e.g., mamastroviruses 6, 8, and 9), require capsid processing by trypsin to infect cells. Several human and primate cell lines, especially those derived from the gastrointestinal tracts, are susceptible to astrovirus ^2^. Recently, astroviruses were shown to replicate in a human intestine-derived organoid ^3^, and HAstV, especially VA, has been detected in the central nervous system (CNS) of immunocompromised patients ^4-11^. There are also cases of viremia in humans ^12,13^; however, the mechanism underlying the viral spread from the intestinal tract or other primary infection sites is unknown.

Astrovirus genomes have three open reading frames (ORFs): ORF1a, ORF1b, and ORF2. ORF1a and ORF1b encode nsP1a and nsP1ab, which become subdivided into a protease, VPg, and RdRp, as well as several proteins of unknown function. ORF2 encodes a capsid protein precursor (VP90), which is cleaved by caspase to form VP70. Extracellular trypsin-mediated processing results in the formation of a mature capsid consisting of the core (VP34) and spike proteins (VP27), as well as the removal of some spike proteins (VP25). Since antibodies that target the spike are neutralizing ^14,15^, VP27 is believed to contain the receptor binding domain. Trypsin treatment remarkably changes HAstV structure ^16^, possibly facilitating receptor interaction.

During cellular infection, HAstV is thought to initially bind to a surface-expressed polysaccharide via the VP27 region that is conserved among serotypes ^17^. It is then thought to enter the cell via clathrin-mediated endocytosis, escape the late endosomes ^18^, and be uncoated by protein disulfide isomerase A4 (PDIA4) to release its genome ^19^. However, a functional receptor for HAstV has not been discovered, and the entry mechanism has not been fully elucidated.

Here, we identified neonatal Fc receptor (FcRn) as a receptor of HAstV using genome-wide CRISPR-Cas9 screening. Although FcRn was directly bound to a spike protein (VP27), depletion of FcRn did not impair HAstV binding to the cell surface. However, HAstV infection was dramatically prevented in FcRn-deficient cells, indicating the existence of other binding factors alongside FcRn in viral uptake. Our findings indicate that FcRn is an important factor for HAstV infection and provide insight into the mechanism underlying HAstV entry.

## Results

### Identification of host genes essential to HAstV infection

We performed CRISPR knockout screenings with type 4 HAstV on Caco2 cells, which are highly susceptible to HAstV infection. HAstV does not normally induce death in Caco2 cells ^20^; however, apoptosis can be induced at higher multiplicities of infection (MOIs) ^21^. The cells were cultured with HAstV in a trypsin-containing medium, and we observed extensive cell death (Fig. S1).

A single-guide RNA (sgRNA) library pool was transduced into Cas9-expressing Caco2 cells. Cells were exposed to HAstV4 and incubated in a trypsin-containing medium for 2 days, with the escaped cells then incubated to increase their numbers (Fig. 1A). We repeated this infection-amplification step five times in the first round and then twice in the second round. Genomic DNA was extracted from the escaped cells, and the sgRNA regions were amplified and sequenced via next-generation sequencing (NGS). The results showed that the genes *FCGRT* and *B2M* could be involved in HAstV susceptibility (Fig. 1B). These genes encode the α (or heavy) chain (encoded by *FCGRT*) and β_2_ microglobulin (β_2_m; encoded by *B2M*) of the neonatal Fc receptor, a non-classical MHC class I molecule.

**Figure 1.**
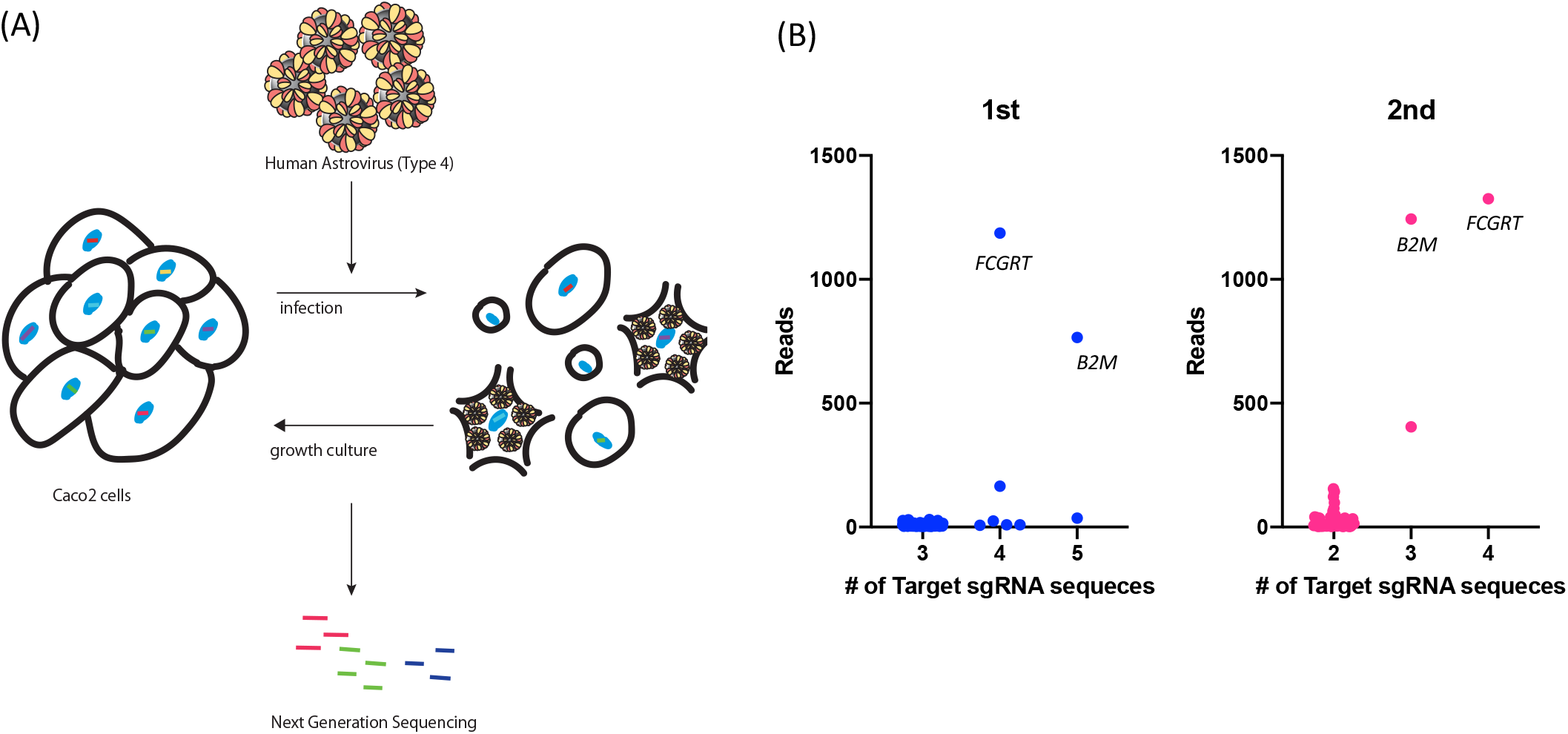
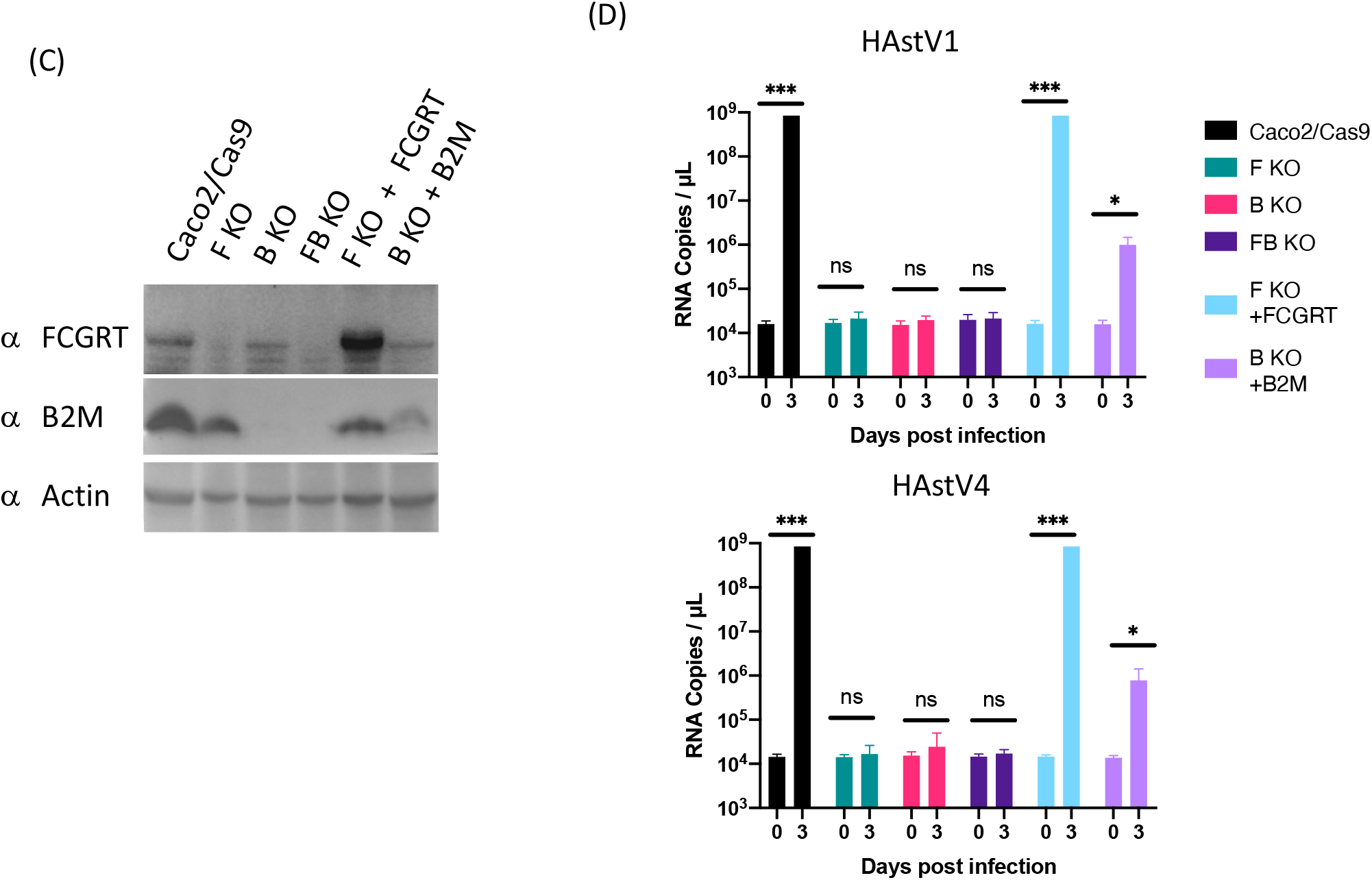
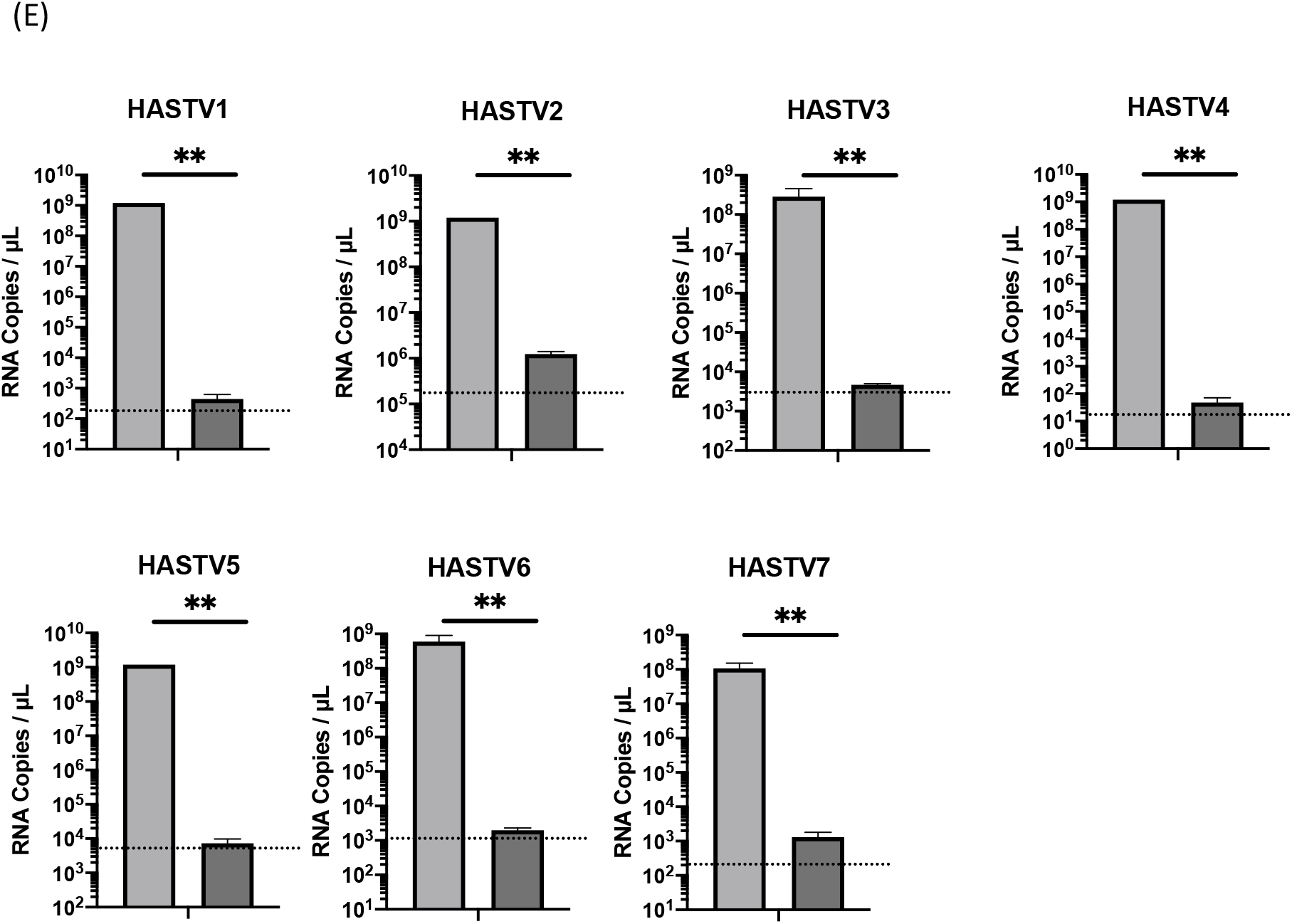
(A) Schema of the screening method for cells that escaped HAstV infection. Caco2 cells transduced with a CRISPR-Cas9 library pool were infected with HAstV and cultured for 2 days; the medium was replaced with a growth medium and inoculated with HAstV 4 again. This cycle was repeated independently several times: five times in the first round and twice in the second round. After the escaped cells were screened, genomic DNA was purified, and sgRNA sequences were determined via next-generation sequencing (NGS). (B) The library included five or six kinds of sgRNA sequences for every gene. The x-axis shows how many different sgRNA sequences per gene were detected via NGS, while the y-axis shows the total read counts per gene. sgRNA of *FCGRT* and *B2M* were commonly observed in the first (blue) and second (pink) rounds. (C) Detection of *FCGRT* and *B2M* in corresponding knockout Caco2 cells. Actin was used as a loading control. (D) Knockout of *FCGRT* or *B2M* inhibited HAstV infection in Caco2 cells. Cells were incubated with HAstV1 or 4 (MOI 0.05) for 1 hour and washed twice, then incubated in a fresh medium containing 5 µg/ml trypsin. Culture supernatants were collected immediately after adding fresh medium (Day 0) and at 3 days post-infection (Day 3). RNA copies in the culture supernatant were determined via qRT-PCR. Each data bar represents the mean for eight wells of inoculated cells. Error bars denote SD. Each experiment was performed three times, and representative data are shown in this figure. Significance was determined by Mann-Whitney test (***, *P* < 0.0005; *, *P* < 0.05); ns, not significant). (E) Knocking out *FCGRT* and *B2M* (FB KO) completely inhibited infection of every type of human astrovirus in Caco2 cells. Normal (light gray) or FB KO Caco2 (dark gray) cells were incubated in a culture medium containing trypsin-activated HAstV for an hour. After washing with the medium three times, the cells were cultured with 5 µg/mL trypsin for 3 days at 37°C. The amount of genomic RNA of each virus was determined by qRT-PCR. Each data bar represents the mean for eight wells of inoculated cells. Error bars denote SD. Each experiment was performed two or more times, and representative data are shown in this figure. Significance was determined by the Mann–Whitney test (**, *P* < 0.005).

To determine the role of these gene products in HAstV infection of Caco2 cells, we established *FCGRT*- or *B2M*-deleted Caco2 cells by CRISPR-Cas9. Since the α chain and β_2_m may form a heterodimer but could also work separately in HAstV infection, we also established *FCGRT*-*B2M* double-knockout (FB KO) Caco2 cells. Deletion of *FCGRT* (F KO) and/or *B2M* (B KO) was confirmed via western blotting (Fig 1C). HAstV4 RNA was greatly amplified in infected parent Caco2/Cas9 cells and showed much less amplification in F KO, B KO, or FB KO Caco2 cells (Fig. 1D). Additionally, HAstV1, which is detected with relatively high frequency, was added to the knockout cells, with the results again showing little amplification in the knockout lines. To confirm that the loss of susceptibility was caused by the gene deletions, each gene was added back to cells by lentivirus encoding sgRNA-resistant *FCGRT* or *B2M*. The addition of *FCGRT* and *B2M* completely and partially restored, respectively, the HAstV RNA level to that in the parent cells (Fig. 1C–D). Additionally, the susceptibilities of FB KO Caco2 cells to other HAstVs 2, 3, 5, 6, and 7 serotypes were analyzed, and results showed similar differences in amplification compared to parent cells. These findings suggest that FcRn is essential in HAstV infection of Caco2 cells.

### Astrovirus susceptibility reduced by FcRn deletion in human intestinal organoid

The primary target of HAstV is intestinal epithelia, and astrovirus infections were recently established in organoids derived from the small intestine or colon ^3^. Accordingly, we investigated whether FcRn was involved in HAstV susceptibility in a human intestinal organoid derived from normal human ileum (Ileum-1). No *FCGRT* or *B2M* expression was observed in FCGRT- or B2M-knockout Ileum-1, respectively (Fig. 2A). HAstV4 activated by trypsin treatment was inoculated into the Ileum-1 monolayer. The increase in HAstV4 RNA level was approximated to that shown in a human enteric organoid ^3^. Ileum-1 was not cultured with trypsin because it was easier to harvest than Caco2 cells; therefore, a newly produced virus could not be activated to infect, resulting in lower HAstV amplification than that seen with the Caco2 cells. HAstV4 RNA was significantly reduced in *FCGRT*- or *B2M*-knockout Ileum-1 (Fig. 3B), suggesting that FcRn is involved in HAstV replication in the intestinal organoid.

**Figure 2.**
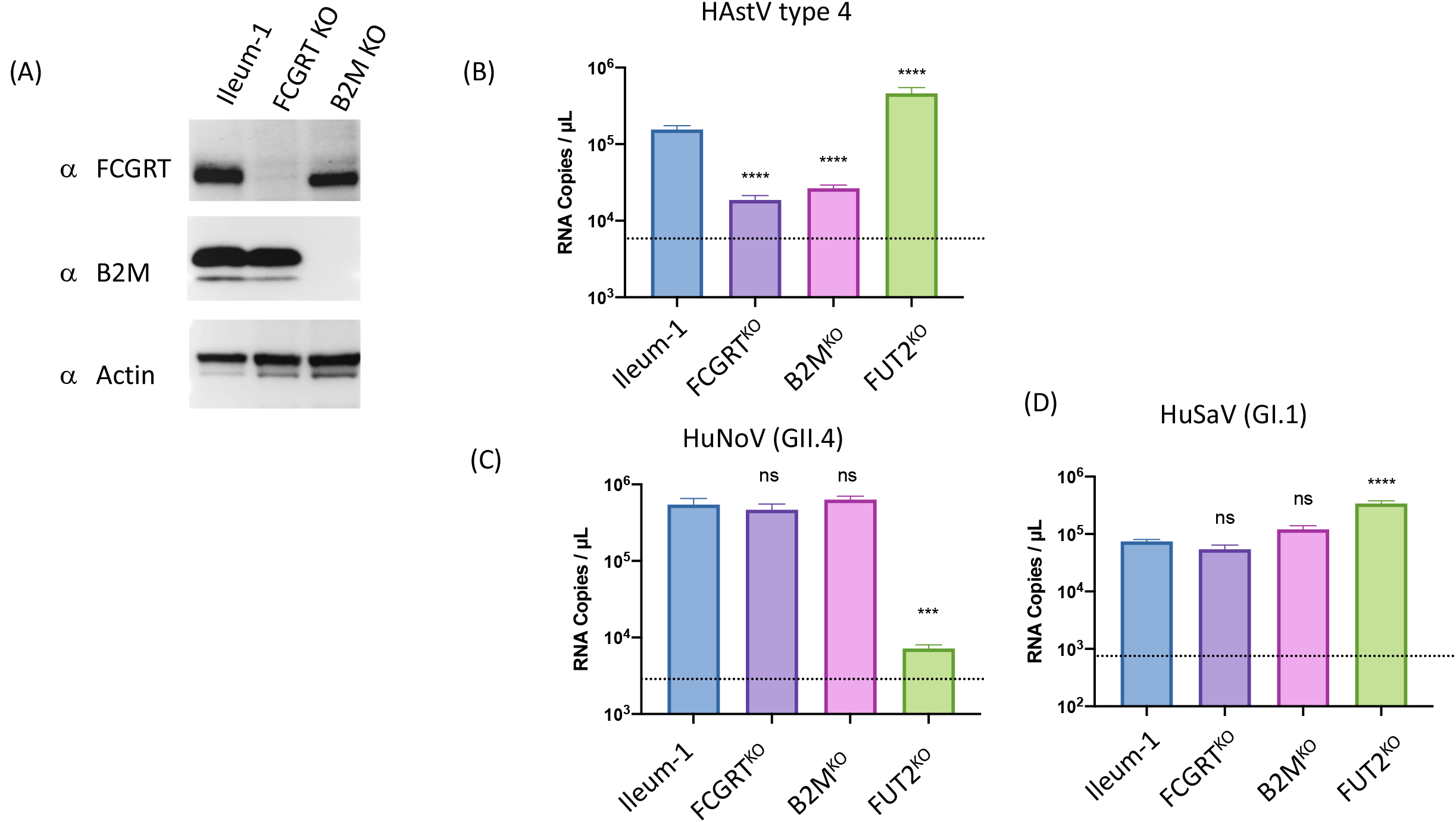
Knocking out *FCGRT* or *B2M* prevents infection of type-4 human astrovirus in an ileum-derived human intestinal organoid (Ileum-1). (A) Detection of *FCGRT* and *B2M* in corresponding knockout Ileum-1 cells. Actin was used as a loading control. (B–D) Each gene-knockout monolayered Ileum-1 was inoculated with a culture medium containing trypsin-activated HAstV4 (B) or stool filtrate containing either GII.4-type HuNoV (C) or GI.1-type HuSaV (D). After washing twice with 3+ medium, cells were cultured at 37°C for 3 days. Genomic RNA copies of each virus were determined by qRT-PCR. Each data bar represents the mean for three wells of inoculated monolayers. Error bars denote SD. Each experiment was performed twice; representative data are shown in this figure. Significance was determined by one-way ANOVA with Dunnett’s multiple comparison test (****, *P* < 0.0001; ns, not significant). The dotted line indicates RNA copies in the culture supernatant at 1 hour post-infection.

**Figure 3.**
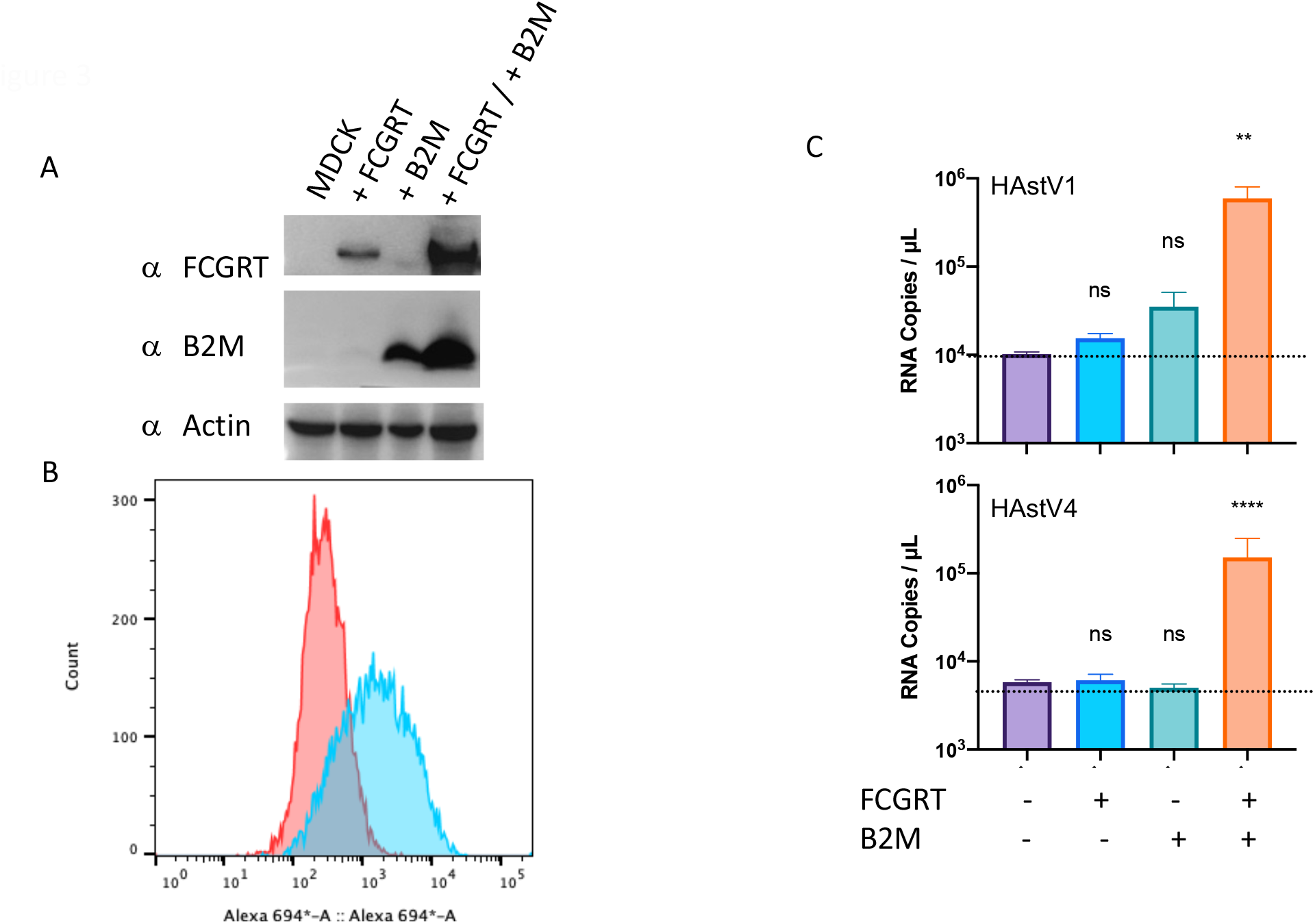
Exogenous co-expression of *FCGRT* and *B2M* induced MDCK cell susceptibility to HAstV. (A) *FCGRT, B2M*, or *FCGRT* and *B2M* were transduced into MDCK cells using a lentiviral vector and stably expressed. *FCGRT* and *B2M* were detected in MDCK cells transduced with each gene. Actin was used as a loading control. (B) Harvested MDCK cells (red) or MDCK cells transduced with *FCGRT* and *B2M* (blue) were treated with rabbit anti-*FCGRT* IgG and subsequently with Alexa 694-conjugated anti-rabbit IgG. The cells were analyzed using a flow cytometer. (C) The MDCK cells were inoculated in a culture medium containing trypsin-activated HAstV for 3 hours. After washing with the medium three times, cells were cultured with 5 µg/mL trypsin at 37°C for 6 days. Genomic RNA copies of each virus were determined by RT-qPCR. Each experiment was performed two times, with five technical replicates in each experiment, and representative data are shown in this figure. Error bars denote SD. Significance was determined by one-way ANOVA with Dunnett’s multiple comparison test (****, *P* < 0.0001; **, *P* < 0.01; ns, not significant). The dotted line indicates RNA copies in the culture supernatant at 3 hours post-infection.

Since enterovirus B also uses FcRn as a receptor^22,23^, the replication of other gastroenteritis viruses—GII.4-type human norovirus (HuNoV) and GI.1-type human sapovirus (HuSaV)—was also analyzed in the knockout Ileum-1. Before evaluating the virus susceptibility genes, we examined whether HuNoV and HuSaV could replicate in Ileum-1. A fecal specimen containing GII.4-type HuNoV was inoculated into Ileum-1, and progeny production in the culture supernatant was determined 3 days post-infection. As shown previously ^24^, deletion of the fucosyltransferase-2 gene *(FUT2)* from Ileum-1 greatly reduced Ulex europaeus agglutinin-1 (UEA-1) binding to the cellular surface, and the susceptibility to GII.4-type HuNoV was abolished (Fig. S2A–B). GI.1-type HuSaV amplified only when sodium glycocholate (GCA), which is essential for HuSaV infection of HuTu80 cells ^25^, was included in the culture medium (Fig. S2C). In contrast to that of HAstV4, *FCGRT* or *B2M* deletion did not change susceptibilities to HuNoV and HuSaV infections (Fig. 3C–D). These findings suggest that FcRn is not universally used in intestinal virus infection, though it is used by enterovirus B and HAstV.

### Exogenous expression of *FCGRT* and *B2M* renders unsusceptible cells permissive to astrovirus

To further assess the role of FcRn in the susceptibility to HAstV, *FCGRT* or *B2M* were transduced into Madin-Darby Canine Kidney (MDCK) cells, which are unsusceptible to HAstV ^2^. Transduction of *FCGRT* or *B2M* into MDCK cells resulted in the expression of their RNA, with co-transduction of *FCGRT* and *B2M* further enhancing this (Fig. 3A), and FCGRT was expressed on the cell surface (Fig. 3B). HAstV1 and HAstV4 RNA were not amplified in *FCGRT*- or *B2M*-transduced MDCK cells; however, cells that expressed both *FCGRT* and *B2M* became permissive to HAstV1 and HAstV4. These results were consistent with those of the Caco2 cells, and both expressions were required for HAstV infection.

### FcRn is involved in the early phase of infection

To investigate which step of the viral life cycle FcRn was engaged in, Caco2 cells were inoculated with activated HAstV and then cultured without trypsin, which prevented infection by newly produced HAstV. First, titers of surface-bound HAstV1 or HAstV4 were compared between Caco2 and MDCK cells that expressed or did not express FcRn: results showed that FcRn expression did not change the bound-virus titers (Fig. 4A), indicating that HAstV binding is not affected by FcRn expression and that other molecule(s) may be involved.

**Figure 4.**
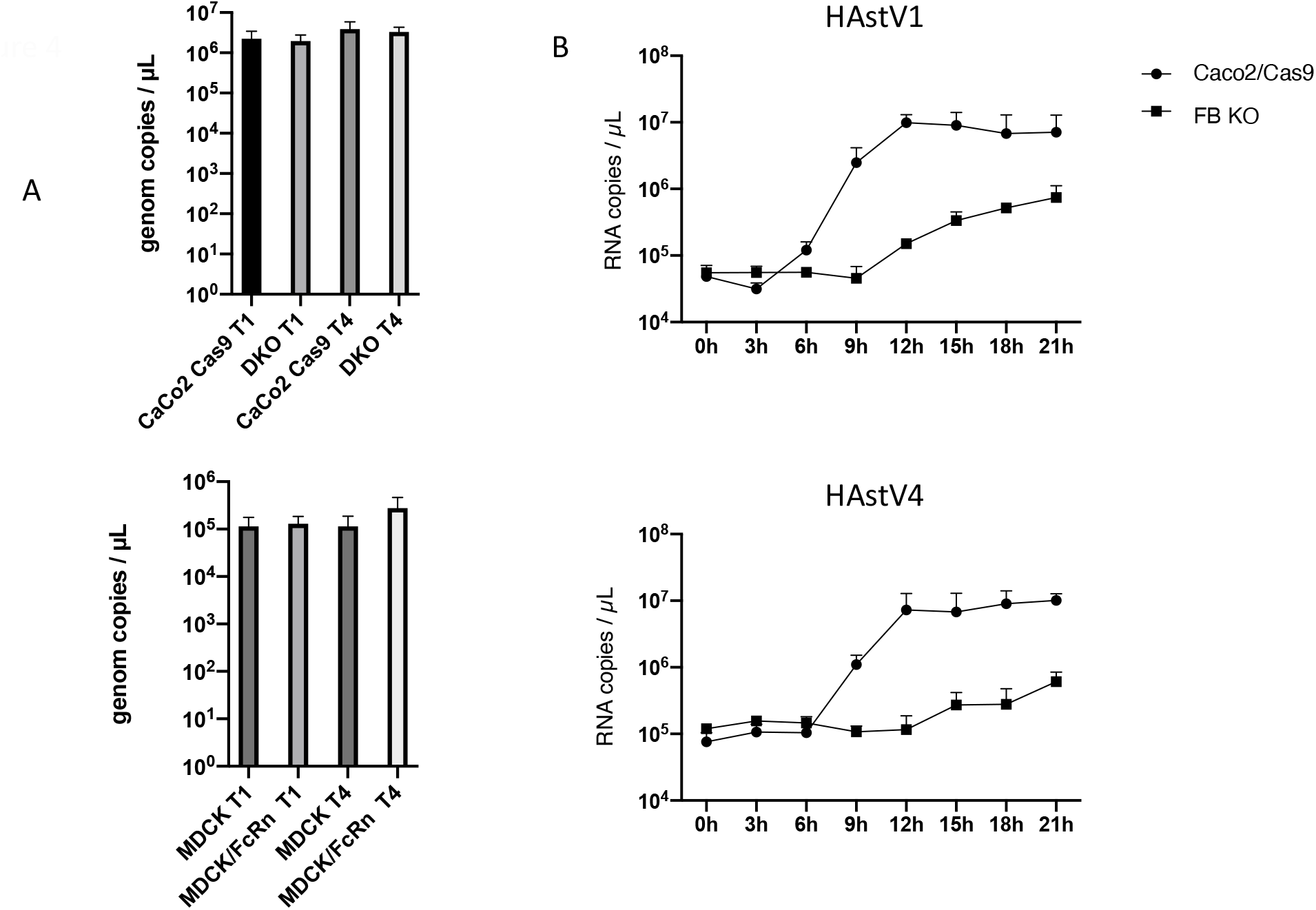
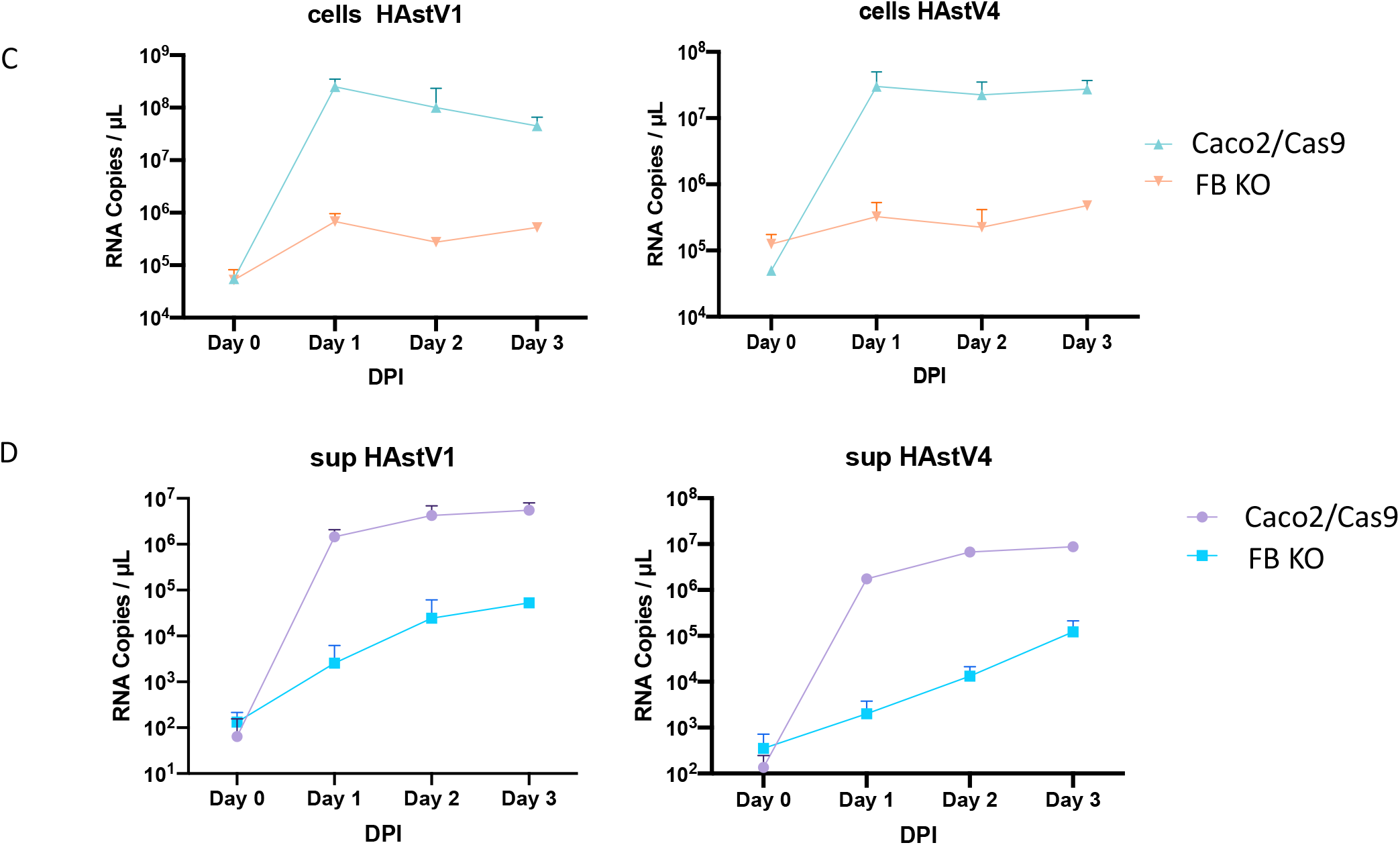
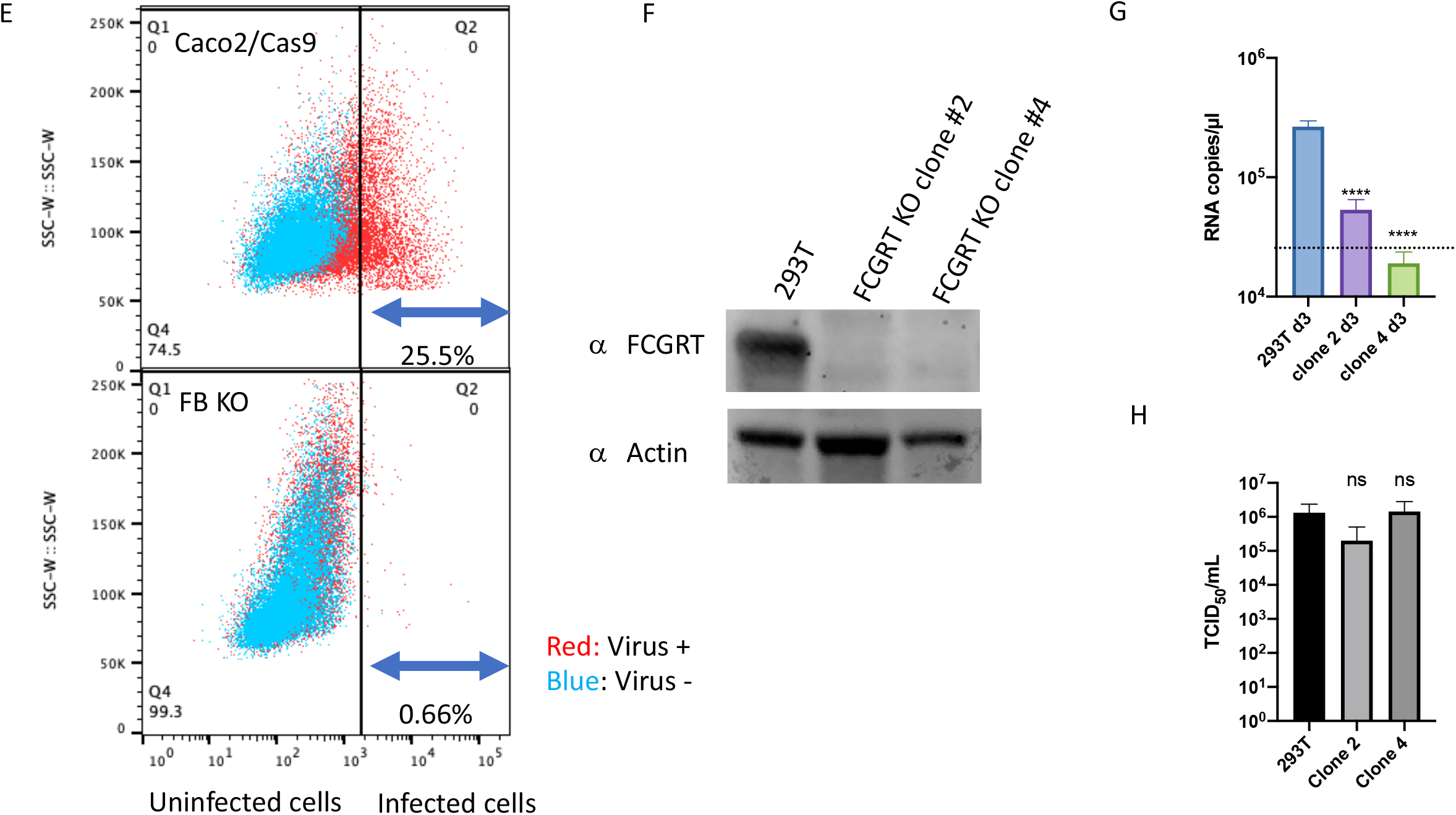
(A) The titer of viruses bound to the cell surface did not change with FcRn status. Caco2 or MDCK cells were incubated with activated HAstV (MOI 50) at 4°C for 1 hour. After washing with MEM (-) three times, RNA was extracted from the cells with the bound virus, and copies were quantified by qRT-PCR. Each data bar represents the mean for four wells. Error bars denote SD. (B) Viral RNA copies of single-step growth in the cells were inhibited in FcRn-KO cells. Trypsin-activated HAstV1 or HAstV4 was inoculated into the cells (MOI 1). At each time point, the culture medium was aspirated from the wells, and the cells were washed with PBS twice. RNA was extracted from cells, and copies were quantified by RT-qPCR. Each point represents the mean for five wells. Error bars denote SD. (C–D) Viral RNA copies in the cells and culture supernatant were inhibited in FcRn-KO cells. Trypsin-activated HAstV1 or 4 was inoculated into the cells (MOI 1). Every day, culture medium and cells (D) were collected separately. The cells were washed with PBS twice, and RNA was extracted. RNA copies in the culture medium and cells were quantified by RT-qPCR. Each data point represents the mean for four wells. Error bars denote SD. (E) Infected cells were detected using PrimeFlow technology. Caco2 cells were exposed to trypsin-activated HAstV4 (MOI 10) and incubated in a trypsin-free medium for 2 days. Cells were detached from the wells with trypsin and treated with a HAstV-specific probe according to manufacturer’s protocol. Probe-hybridized cells were analyzed by Melody (BD) and FLOWJO. Red dots represent HAstV4-infected cells, and blue dots represent uninfected cells. (F–H) Knocking out *FCGRT* prevents HAstV4 infection in 293T cells. (F) Detection of *FCGRT* in 293T cells and *FCGRT*-KO clones. Actin was used as a loading control. (G) HAstV4 was incubated with trypsin to activate it. After activation, viruses were treated with FBS to inactivate trypsin and were inoculated into cells. After 1-hour incubation at 37°C and subsequent washing with the medium twice, the cells were cultured at 37°C for 3 days. Genomic RNA copies of each virus were determined by RT-qPCR. Each data bar represents the mean for three wells of inoculated monolayers. Error bars denote SEM. Each experiment was performed two times, and representative data are shown in this figure. Significance was determined by one-way ANOVA with Dunnett’s multiple comparison test (****, P < 0.0001; *, P < 0.05; ns, not significant). The dotted line represents RNA copies in culture supernatant at 1 hour post-infection. (H) Viral RNA extracted from HAstV4 was transfected into 293T cells and *FCGRT*-KO clones. After 72 hours of incubation, the culture supernatant was serially diluted and inoculated into Caco2 cells. After 5 days of incubation, Median Tissue Culture Infectious Dose (TCID50) was determined. Each data bar represents the mean for four wells of transfected 293T cells or KO clones. Each experiment was performed three times, and representative data are shown in this figure. Error bars denote SD. Significance was determined by one-way ANOVA with Dunnett’s multiple comparison test (ns, not significant).

Second, when observed over 21 hours post-infection, HAstV RNA levels plateaued at 12 hours in parent Caco2 cells but were still gradually increasing in FB KO Caco2 cells (Fig. 4B). We then extended the observation period to 3 days post-infection and found that viral RNA levels plateaued on day 1 post-infection in KO cells (Fig. 4C). Progeny production in Caco2 cells increased by approximately 1 × 10^4^ times that produced on the first day post-infection. Viral RNA levels in KO Caco2 cells increased to a lesser extent than that observed in the previous results shown in Fig. 1E, presumably due to the titer at inoculation in that experiment being 20 times higher (Fig. 4D).

Third, we determined the percentage of infected cells by detecting RNA fragments in the cells using the PrimeFlow assay kit (Thermo), which uses probes that hybridize to subgenomic RNA regions and highlight infected cells. The cells used had been previously infected with HAstV4 at an MOI of 10. Infected parent Caco2 cells shifted right, with 25.5 % containing highly replicated genomic or subgenomic RNA (Fig. 4E); meanwhile, only a few KO cells were observed to possess viral RNA (0.66%), indicating that few KO cells were infected and produced progeny virus (Fig. 4E).

Lastly, since FcRn can mediate the endocytosis of antibodies as well as releasing them ^26,27^, we examined whether FcRn was involved in viral shedding. Since transfection of viral RNA into cells can yield infectious virion ^28^, we extracted HAstV4 genomic RNA and transfected it into 293T or 293T/FCGRT^KO^ cells, and the amount of infectious virion was compared with that of infected Caco2 cells. 293T cells have a slight susceptibility to HAstV^2^, and *FCGRT* depletion abolished this (Fig. 4F–G). There was no significant difference in the levels of infectious virion shedding in each cell line (Fig. 4H). In conclusion, although there may be FcRn-independent infection machinery, FcRn is involved in the early stages of viral replication.

### FcRn directly binds to human astrovirus

To investigate whether FcRn directly binds to HAstV, we treated Caco2 cells with an anti-FCGRT antibody before virus inoculation to block the interaction between FcRn and HAstV. The results showed that anti-FCGRT antibody dose-dependently reduced amplification (Fig. 5A). An enzyme-linked immunosorbent assay (ELISA) was then performed to examine the direct interaction between recombinant FcRn protein and HAstV1 or HAstV4. Immobilized recombinant FcRn was treated with serially diluted HAstV1 or 4, then detected using an anti-astrovirus antibody. The assay revealed dose-dependent binding of HAstV1 and, to a lesser extent, HAstV4 (Fig. 5B). Since no difference was observed in the sensitivity of the anti-astrovirus antibody for each type of HAstV (Fig. 5C), there could be a difference in the binding affinity of the serotypes to FcRn. Additionally, we showed that recombinant FcRn was directly bound to the recombinant spike protein (VP27) derived from HAstV types 1, 4, and 7, which was expressed by *Escherichia coli*. Although the amount of immobilized VP27s was not different between serotypes, VP27 of HAstV7 showed the highest affinity to recombinant FcRn, followed by that of HastV1, and that of HAstV4 the lowest. These findings suggest that FcRn directly interacts with the spike of HAstV and works as a functional receptor of HAstV.

**Figure 5.**
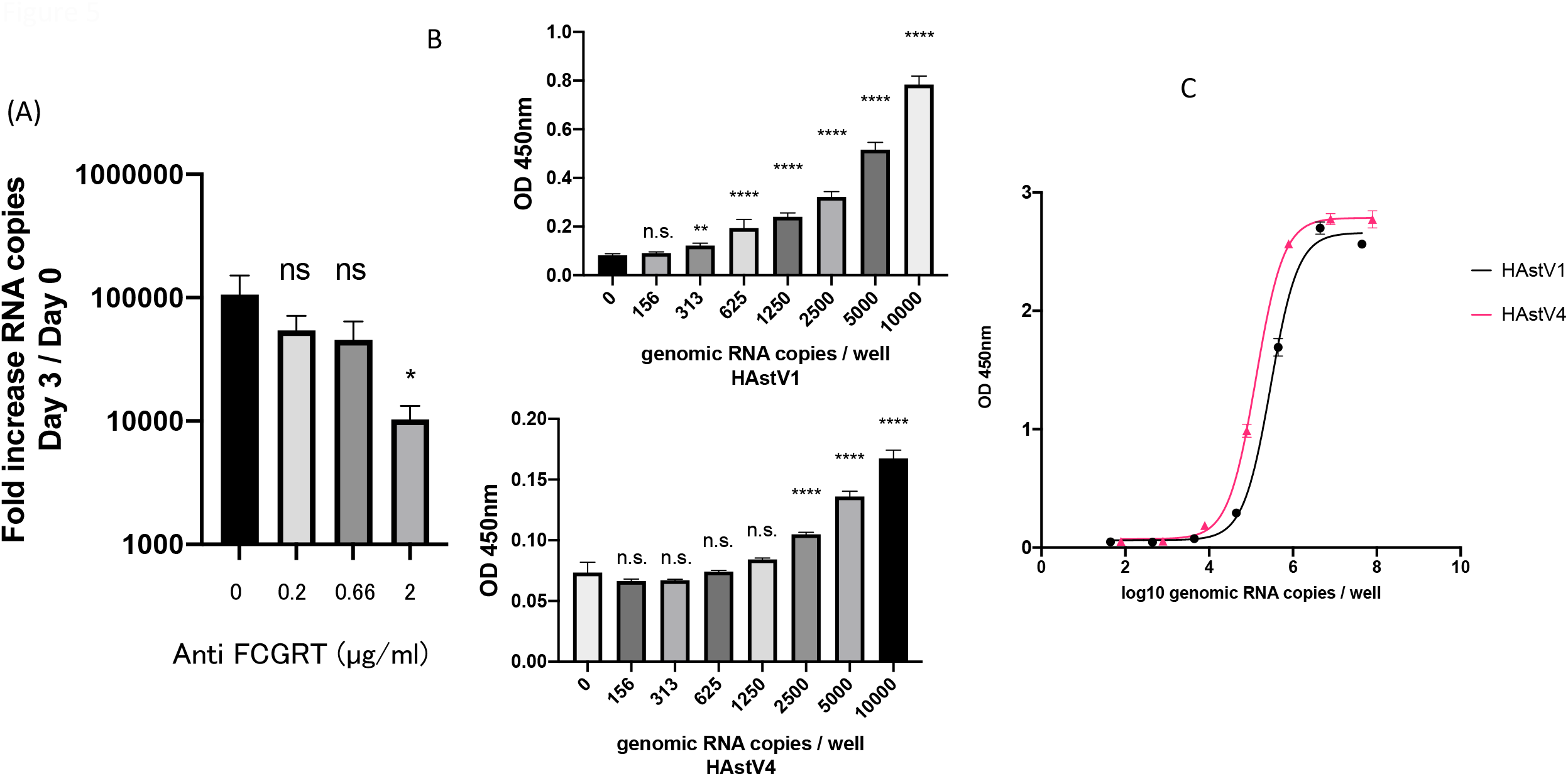
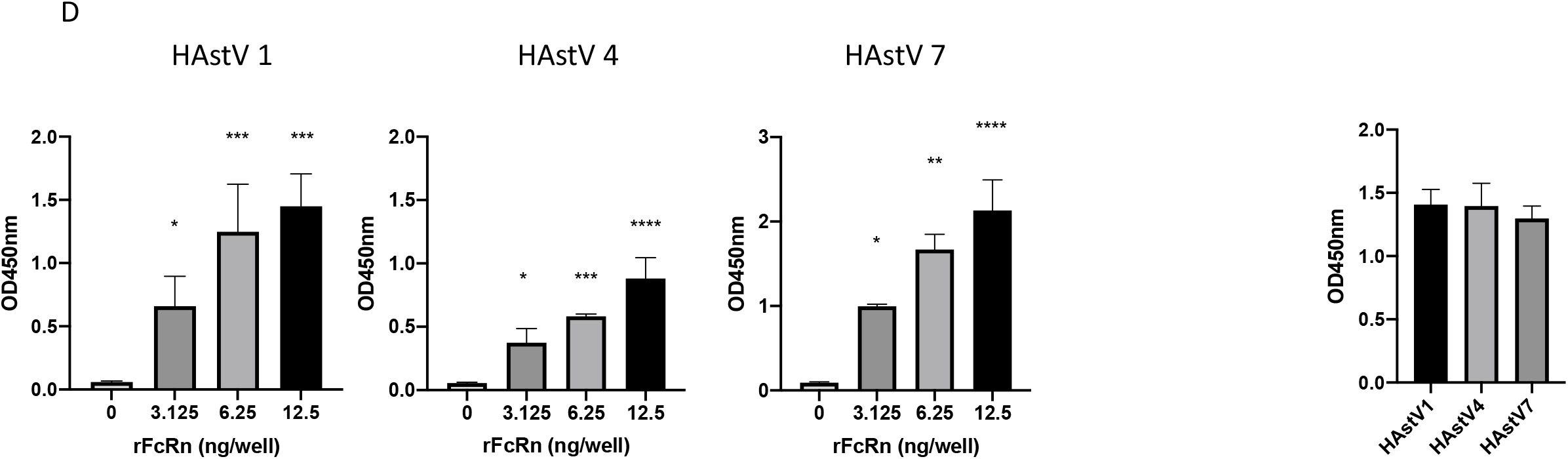
(A) Anti-*FCGRT* antibodies block HAstV1 infection. Caco2 cells were cultured with 2, 0.66, or 0.2 µg/mL of anti-*FCGRT* antibody (NBP1-89128: NOVUS) for 1 hour before infection. The Caco2 cells were incubated with the activated virus (MOI 0.005) and anti-*FCGRT* at the indicated concentrations for 1 hour. The cells were then washed with fresh medium three times and cultured for 3 days with anti-*FCGRT* at the indicated concentrations. Error bars are mean ± SEM (n = 3). Significance was determined by one-way ANOVA with Dunnett’s multiple comparison test (*, *P* < 0.05; ns, not significant). (B) ELISA measuring the binding affinity of FcRn and HAstV1 or 4. Immobilized recombinant FcRn was incubated with serial dilutions of each HAstV. HAstV captured by FcRn was detected by an anti-astrovirus antibody (8E7). Error bars are mean ± SEM (n = 10). Data are representative of two independent experiments. Significance was determined by one-way ANOVA with Dunnett’s multiple comparison test (****, *P* < 0.0001; **, *P* < 0.01; ns, not significant). (C) Direct antigen ELISA measuring the binding affinity of anti-HAstV to immobilized purified HAstV. Each immobilized HAstV was detected by an anti-astrovirus antibody (8E7). Error bars are mean ± SEM (n = 5). (D) Each type of His-tagged VP27 was expressed by BL21. Purified His-tagged VP27 (2.5 µg) was immobilized and incubated with diluted recombinant FcRn. FcRn bound to VP27 was detected by an anti-FcRn antibody. Data are representative of two independent experiments. Error bars are mean ± SEM (n = 3). Significance was determined by one-way ANOVA with Dunnett’s multiple comparison test (****, *P* < 0.0001; ***, *P* < 0.001; **, *P* < 0.01; *, *P* < 0.05). Each type of immobilized His-tagged VP27 was detected with HRP-conjugated anti-His antibody (MBL), as shown in the right panel.

## Discussion

Here, we showed that FcRn depletion in Caco2 cells and a human intestinal organoid inhibited human astrovirus infection and that exogenous FcRn expression rendered MDCK cells susceptible to HAstV. Meanwhile, FcRn had no role in GII.4 human norovirus and GI.1 human sapovirus infection. We also showed that FcRn was directly bound to the HAstV spike protein, indicating that FcRn functioned as a receptor for HAstV.

FcRn is categorized as a non-classical MHC class I molecule. It is highly expressed in infants’ intestines, where it functions to transfer maternal IgG from the milk in the intestines ^29,30^. IgG binds FcRn at the apical membrane of the intestines and is then released at the basolateral membrane in a process regulated based on the pH ^26^. FcRn is expressed not only in the intestinal epithelium of neonatal rodents but also ubiquitously in various tissues of adults ^27^.

FcRn is expressed in the epithelial cells of the intestinal tract, kidney, placenta, and liver, as well as in hematopoietic cells, vascular endothelia, and at the blood-brain barrier (BBB), and works to recycle IgGs or albumin by transporting them into and then out of cells ^27^. The primary target of HAstV is the intestines, and infection can cause gastroenteritis and, less frequently, encephalitis in immunocompromised patients. VA1 (mamastrovirus 9) ^4,6,8,10,11^ is the most frequently detected, while MLB (mamastrovirus 6) ^5^ and HAstV1 and 4 (mamastrovirus 1) ^9,12^ have also been detected in immunocompromised patients. FcRn is bound to the spike protein (VP27) of HAstVs (Fig. 5D); however, the amino acid sequences of the spike regions of HAstV1 and VA1 are quite different ^20^ and, therefore, VA1 and MLB could not use FcRn as a receptor. Indeed, while VA1 infected primary astrocytes and immortalized neuronal cell lines, a slight HAstV4 infection was observed ^31^. FcRn is expressed in vascular endothelial cells in the brain as well as some neuronal cell lines, according to the Human Protein Atlas ^32^, indicating that HAstV infecting the intestines may escape host immunity and transit through the bloodstream to infect the CNS. Further investigation is required to find out if HAstVs infect neuronal cells via FcRn and if novel-type HAstVs (VA and MLB) use FcRn as a receptor.

Cleavage of capsid proteins by trypsin, which induces efficient HAstV infection, exposes the spike protein VP27 and allows neutralizing monoclonal antibodies against HAstV to bind to VP27 ^14 15^. Neutralizing antibodies could block the binding of VP27 with FcRn because we showed that FcRn directly binds to VP27. After their binding, HAstV could be taken into the cell via the clathrin heavy chain (CHC) pathway; it was shown that CHC inhibitor or CHC siRNA reduced HAstV8 infectivity ^18^ and FcRn-mediated transcytosis of IgG was also associated with clathrin ^26^. Taken together, HAstVs bind to surface-expressed FcRn and enters the cell. Then, after entry, FcRn itself may have a role in viral uncoating because FcRn functions pH-dependently as an uncoating receptor for enterovirus B ^22^. Moreover, PDIA4 was reported as an uncoating molecule for HAstVs 1 and 8 ^19^ but not for HAstV 2, which may cause the difference in susceptibility between these. Our experiments on FcRn-KO Caco2 cells infected with higher titers (Fig. 4D) also suggest that independent FcRn-mediated infectious routes exist. In contrast, although FcRn has the role of releasing IgG or albumin from the cytoplasm via transcytotic activity in genital tract epithelial cells for human immunodeficiency virus-1 ^33^, FcRn was not associated with HAstV shedding. Thus, our findings provide new insights into understanding the mechanism underlying FcRn-mediated HAstV entry.

## Supporting information

Suulement figures

## Acknowledgments

This work is supported in part by the Japan Agency for Medical Research and Development (AMED) 17fk0108034h0501and 18fk0108034h0502 (A.N., K.K.) and a Grant-in-Aid for Scientific Research (C) (K.H).

## Author contributions

K.H., A.N.,and K.K. designed research; K.H., R.T.-T., A.K., and A.N. performed research; K.H., and R.T.-T. analyzed data; and K.H. wrote the paper.

## STAR Methods

### Cells and viruses

Caco2 cells (ATCC) were cultured in MEM (Minimal Essential Medium; Nacalai tesque) containing 10% FBS, 1% nonessential amino acids (Nacalai Tesque, 06344-56), 1% sodium pyruvate (Gibco, 11360-070), and 1% antibiotic-antimycotic (Gibco, 15240-062). MDCK cells (ATCC) were cultured in Dulbecco’s minimal essential medium (Nacalai Tesque) containing 10% FBS and 1× penicillin-streptomycin (Gibco, 15140-122). Intestinal organoid (Ileum-1) was isolated at Keio University of Medicine and cultured in IntestiCult Organoid Growth Medium (VERITAS). HAstV1 (VR-1936™), HAstV2 (VR-1943™), HAstV3 (VR-1944™), HAstV5 (VR-1947™), HAstV6 (VR-1948™), and HAstV7 (VR-1949™) were provided by ATCC, and HAstV4 was kindly gifted from Dr. Mitsuaki Oseto^34^.

### Astrovirus infection and quantitative real-time PCR

Astrovirus was treated with 10 µg/ml of trypsin type IX-S (Trypsin from porcine pancreas Type Ⅸ-S: SIGMA, T0303-1G) at 37°C for 1 hour. Caco2 cells cultured on a collagen I-coated 96-well plate (Corning) were inoculated with the virus (MOI 0.05) and incubated at 37°C for 1 hour. After washing with MEM containing 1% nonessential amino acids, 1% sodium pyruvate, 20-mM HEPES (pH 7.5), and 1% antibiotic-antimycotic solution (MEM (-)), the cells were incubated in MEM (-) containing 5 µg/ml Trypsin IX-S for 15 minutes, and the supernatant was collected and labeled as “Day 0.” Viral RNA in the supernatant was quantified using Norovirus G1/G2 high-speed probe detection kit (TOYOBO) with the primers and probe changed for astrovirus. The forward primer sequence was 5ʹ-ACTGCDAAGCAGCTTCGTGA-3ʹ, and the reverse primer sequence was 5ʹ-CTTGCTAGCCATCRCACTTCT-3ʹ. The probe sequence was 5ʹ-ROX-CACWGAAGAGCAACTCCATCGCAT-BHQ2-3ʹ.

### Screening of astrovirus susceptibility genes

Cas9-expressing lentiviral vector (lentiCas9-Blast) and Human GeCKOv2 CRISPR knockout pooled library^35^ were provided by Addgene (# 1000000049). Cas9 was induced in Caco2 cells and was selected with 5 µg/ml of blastcidin-S (TAKARA). Then the GeCKO2 library was induced in the Caco2/Cas9 cells and selected with 5 µg/ml of puromycin (TAKARA). Transduced cells were incubated with MEM (-) plus type 4 HAstV (MOI 5). After 2 days, the culture medium was replaced with MEM (+), and the cells were incubated for 6 hours, washed three times with PBS, and then incubated with MEM (-) plus HAstV. These infection-growth cycles were repeated until no surviving cells were visible in Caco2/Cas9 cells. The library-induced Caco2/Cas9 cells, which escaped HAstV infection, were grown in MEM (+), and genomic DNA was then extracted. Integration of the sgRNA sequence into genomic DNA was identified using PCR with the following primers: 5ʹ-GACTATCATATGCTTACCGTAAC-3ʹ and 5ʹ-AAAAAGCACCGACTCGGTGCCAC-3ʹ. Amplified fragments were sequenced via NGS as previously described ^36^.

### Lentiviral vector expressing sgRNA of *FCGRT* or *B2M*

The sequence of the *FCGRT*-specific sgRNA was targeted exon 3 (5’-ACCGCCAAGTTCGCCCTGAA-3’) was cloned into lentiguide-puro vector (Addgene #52963) according to the manufacturer’s protocol. The sequence of the *B2M*-specific sgRNA was targeted to exon 1 (5ʹ-ACTCACGCTGGATAGCCTCC-3ʹ) and cloned into lentiguide-hyg vector, replacing a hygromycin-resistant gene into lentiguide-puro vector.

### Establishing Ileum-1 KO line

Ileum-1 was isolated from ileum. Cas9 and sgRNA were transduced into Ileum-1 by a lentiviral vector, and transduced cells were selected by 2 µg/mL of blastcidine and 2.5 µg/mL of puromycine. After selection, knockout cells were isolated via single-cell cloning.

### Quantification of virus-binding on the cell surface and viral RNA replication in the cells

For quantification of the bound virus, cells were incubated with activated HAstV (MOI 50) at 4°C for 1 hour. After washing with MEM (-) three times, RNA was extracted by Nucleospin 8 (MACHEREY-NAGEL). To quantify viral RNA replication in the cells, HAstV was first treated with trypsin at 37°C for 1 hour to activate it, with trypsin then inactivated by adding FBS, and HastV was then inoculated into cells (MOI 1). At each time point, the supernatant and cells were collected separately. The cells were washed with PBS after aspirating the supernatant. RNA from the cells was extracted using Nucleospin 8 (MACHEREY-NAGEL).

### ELISA for binding assay of FcRn and astrovirus

Recombinant FcRn (Sino Biological) at 12.5 µg/ml was immobilized on a 384-well plate. After blocking with 1% BSA/PBS (bovine serum albumin: SIGMA, A7030-10G), three-times diluted HAstV1 or HAstV4 were incubated on the plate for 2 hours. After washing, wells were treated with anti-astrovirus (8E7) mouse monoclonal IgG (Santa Cruz Biotechnology, sc-53559) as a primary antibody and subsequent HRP-conjugated anti-mouse IgG antibody as a secondary antibody. ELISA POD Substrate TMB Solution (Nacalai Tesque, 05299-54) was used as a substrate, and the absorbance at 450 nm was detected using Ensight (PerkinElmer).

### Recombinant VP27 expression

Each cDNA corresponding to VP27 (residues 394 to 648 of HAstV1 or HAstV4 and 395 to 648 of HAstV7) was cloned with 10× His tag into the pET-52b (Novagen) vector. The expression vector was transformed into BL21 (DE3) (New England Biolabs). The recombinant VP27 was induced with 1-mM IPTG at 37°C for 3 hours after reaching an optical density of 0.5. The induced cells were lysed by ultrasonication in PBS, and VP27 was solubilized by 8 M urea and then purified from soluble lysates by HisTrap metal-affinity chromatography (cytiva).

### ELISA for binding assay of FcRn and recombinant VP27

Diluted recombinant VP27 to 50 µg/ml was immobilized on a 96-well plate. After blocking with 1% BSA/PBS (Bovine Serum Albumin: SIGMA, A7030-10G), recombinant FcRn (Sino Biological, CT071-H27H) was incubated on the plate at 37℃ for 1 hour. After washing, the wells were treated with anti-FcRn mouse monoclonal IgG (Abcam, ab228975) as the first antibody and subsequently the HRP-conjugated anti-mouse IgG antibody (cell signaling, #7074) as a secondary antibody. KPL SureBlue Reserve TMB Microwell Peroxidase Substrate (sera care, 10569771) was used as a substrate, and the absorbance at450 nm was detected usingEnsight (PerkinElmer). Each type of immobilized VP27 was directly detected by HRP-conjugated anti-His antibody (MLB, D291-7).

### Exogeneous expression of *FCGRT* and *B2M*

cDNA of *FCGRT* or *B2M* was cloned into the lentivirus vector-encoded plasmids pLVSIN-puro or pLVSIN-neo vector, respectively. Lentivirus vectors were transduced into MDCK cells. The transduced cells were selected using 2 µg/ml puromycin or 400 µg/ml G-418. CRISPR-resistant mutation for *FCGRT* (5ʹ-ACGGCGAAATTTGCGCTCAA-3ʹ) was introduced in 5ʹ-ACCGCCAAGTTCGCCCTGAA-3ʹ and mutation for *B2M* (5ʹ-ACTCACGTTGTATGGCTTCC-3ʹ) in 5ʹ-ACTCACGCTGGATAGCCTCC-3ʹ. The mutants were transduced in each expression vector (pLVSIN-puro-FCGRT or pLVSIN-neo-B2M) by mutagenesis using prime star max (TAKARA). The selection marker of the resistant *FCGRT* vector was replaced with hygromycin.

### Lentivirus preparation

Lentivirus was prepared as previously described ^36^. Briefly, lentivirus vectors were produced in HEK293T cells by transfecting three component plasmids (i.e., pMDLg/pRRE, pMD2.G, and pRSV-Rev) and pLVSIN- or plentiguide-plasmid using Polyethylenimine “Max” (Polysciences) as a transfection reagent.

### PrimeFlow

PrimeFlow analysis (ThermoFisher Scientific) was performed according to the manufacturer’s protocol. Caco2 cells were infected with HAstV4 (MOI 10) and collected 2 days after infection. HAstV4-specific probes were designed for subgenomic regions of HAstV4. Probe-hybridized cells were analyzed by Melody (BD) and FlowJo software.

### Western blotting

Whole-cell lysates were prepared by adding Laemmli’s SDS/PAGE sample buffer after washing the cells with PBS (−) solution. The lysates were separated by 5–20% polyacrylamide gel (ePAGEL; ATTO) and transferred onto a PVDF membrane using Trans blot turbo (Bio-Rad). The anti-FcRn/*FCGRT* antibody (Novus Biologicals, NBP1-89128), anti-β_2_-microglobulin antibody (Abcam, ab75853), and anti-actin antibody (Santa Cruz Biotechnology, sc-1616) were used as primary antibodies. HRP-conjugated anti-rabbit IgG (Cell Signaling Technology, 7074S) and anti-goat IgG (Santa Cruz Biotechnology, sc-2020) were used as secondary antibodies. The ChemiDoc touch (Bio-Rad) was used to detect proteins visualized by Chemi-Lumi One L Solution A and Chemi-Lumi One L Solution B (Nacalai Tesque, 07880-54).

### Flow cytometric analysis

The expression of *FCGRT* was analyzed using FACS. Cells were trypsinized to detach from the plate. The anti-FcRn/*FCGRT* antibody (Novus Biologicals, NBP1-89128) was incubated with the cells for 30 min on ice. Then, the cells were washed and incubated with Alexa Flour 647 donkey anti-rabbit IgG (ThermoFisher Scientific) for 30 min on ice. Afterward, the cells were washed and subjected to analysis. The antibodies were diluted in Block Ace solution (Waken B Tech), and the washing steps were carried out in ice-cold phosphate-buffered saline without magnesium and calcium salts (PBS (−)). The cells were analyzed using Melody (BD) and the FlowJo software.

### Viral RNA transfection

Genomic RNA of HAstV was extracted from HAstV4 that had been cultured with Caco2 cells under trypsin-containing conditions using a collagen-I coated T75 flask (IWAKI). The virus in the culture supernatant was precipitated by ultracentrifuge, and genomic RNA was purified using the High Pure Viral RNA Kit (Roche) according to the manufacturer’s protocol. 293T or *FCGRT*-KO 293T cells were plated on a 12-well plate a day before transfection, and the culture medium was replaced with DMEM without FBS just before transfection. The cells were transfected with 100 ng of purified RNA and cultured for 48 hours. The culture supernatants were concentrated by ultracentrifugation and serially diluted to determine TCID50 using Caco2 cells.

### Lead contact

Further information and requests for resources and reagents should be directed to and will be fulfilled by the lead contact, Kazuhiko Katayama or Kei Haga.

