## Supplementary figures and images for "Neonatal Fc receptor is a functional receptor for human astrovirus"

### Suulement figures

S-Fig.1

(A)

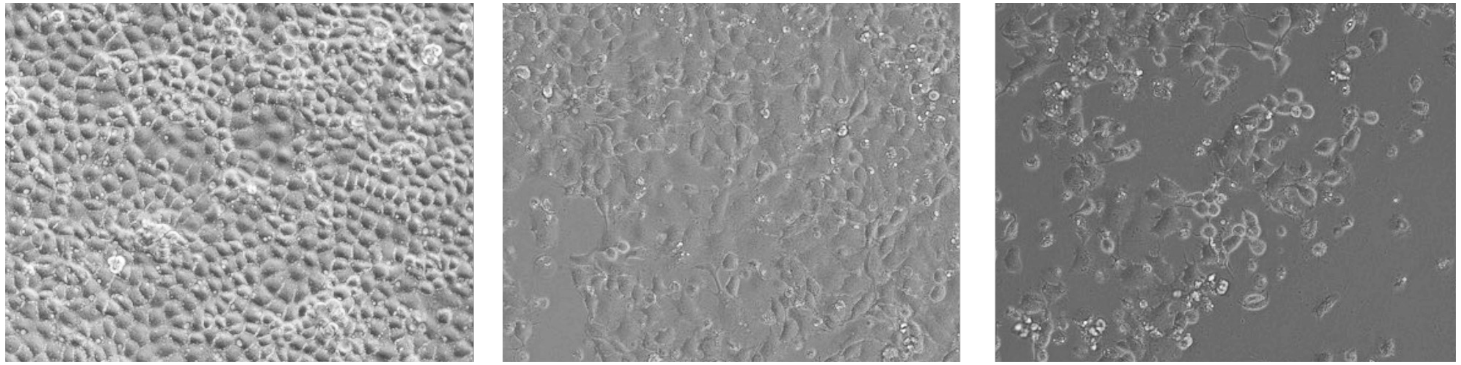

Day 0 3 6

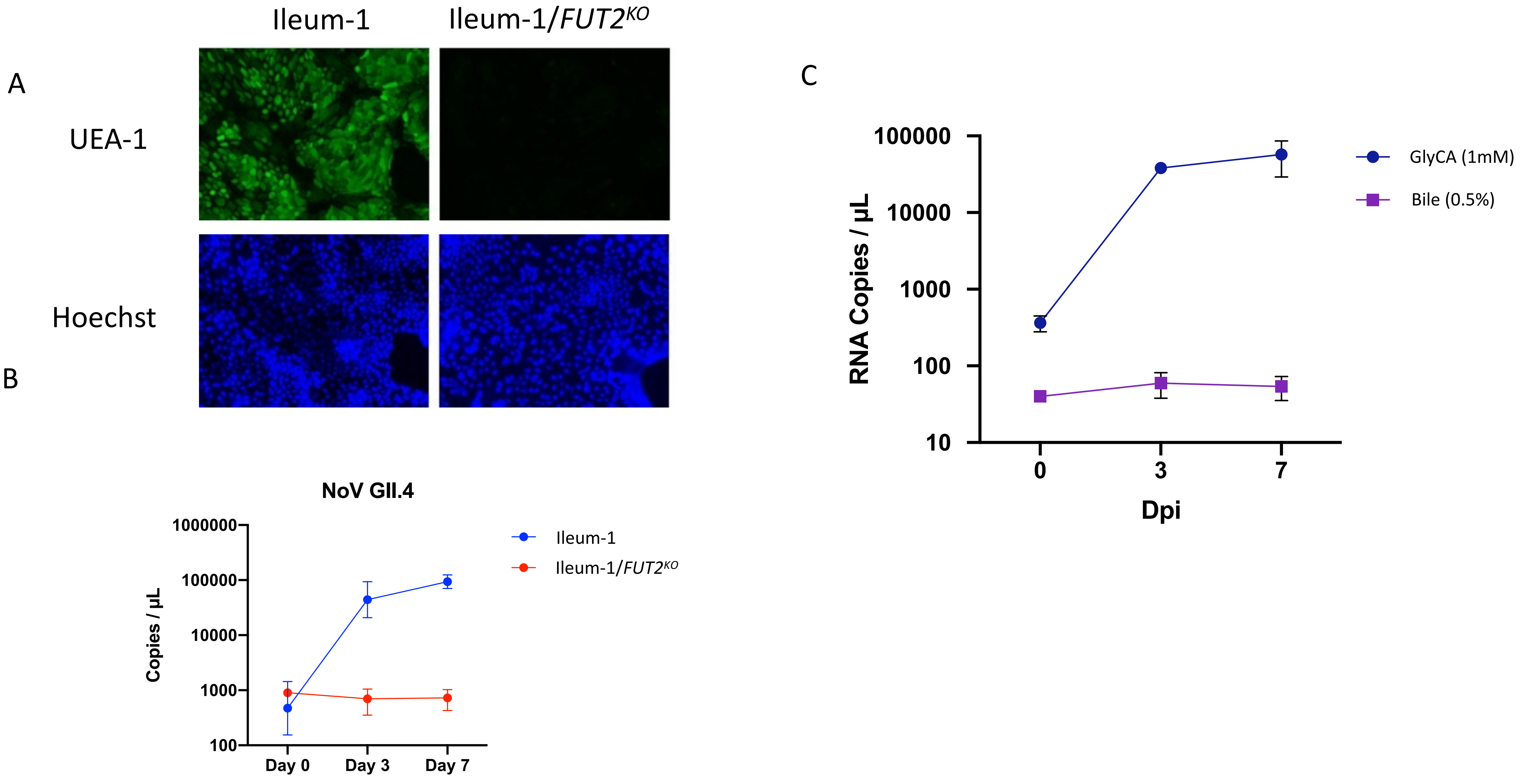
